# Experimental and analytical framework for “mix-and-read” assays based on split luciferase

**DOI:** 10.1101/2022.02.13.480265

**Authors:** Nikki McArthur, Carlos Cruz-Teran, Apoorva Thatavarty, Gregory T. Reeves, Balaji M. Rao

**Author notes:** Address correspondence to, Gregory T. Reeves and Balaji M. Rao.

## Abstract

The use of immunodetection assays including the widely used enzyme-linked immunosorbent assay (ELISA) in applications such as point-of-care detection is often limited by the need for protein immobilization and multiple binding and washing steps. Here, we describe an experimental and analytical framework for the development of simple and modular “mix-and-read” enzymatic complementation assays based on split luciferase that enable sensitive detection and quantification of analytes in solution. In this assay, two engineered protein binders targeting non-overlapping epitopes on the target analyte were each fused to non-active fragments of luciferase to create biosensor probes. Binding proteins to two model targets, lysozyme and Sso6904, were isolated from a combinatorial library of Sso7d mutants using yeast surface display. In the presence of the analyte, probes were brought into close proximity, reconstituting enzymatic activity of luciferase and enabling detection of low picomolar concentrations of the analyte by chemiluminescence. Subsequently, we constructed an equilibrium binding model that relates binding affinities of the binding proteins for the target, assay parameters such as the concentrations of probes used, and assay performance (limit of detection and concentration range over which the target can be quantified). Overall, our experimental and analytical framework provide the foundation for the development of split luciferase assays for detection and quantification of various targets.

## Introduction

Mix-and-read assays, also known as homogeneous immunoassays, are simple analytical techniques used to detect analytes in complex biological fluids. Although enzyme-linked immunosorbent assay (ELISA) is the most widely used assay for analyte detection due to its high sensitivity, it may not be suitable for direct analysis of samples in complex biological fluids or on-site or point-of-care detection.^1–3^ Furthermore, ELISA is a time consuming and complex assay because it requires immobilization of antigens or antibodies on a suitable substrate, multiple washing steps, and long incubation times.^4^

To overcome limitations of ELISA, different types of simple homogeneous phase immunoassays that require no washing steps or protein immobilization have been developed. As shown in **Fig. 1**, the premise of these techniques is that when two sensing components are brought into proximity by the presence of an analyte, a measurable signal, such as luminescence or fluorescence, is generated. For instance, homogeneous open sandwich ELISA has been developed to detect analytes in solution by Förster resonance energy transfer (FRET)^5^, bioluminescence resonance energy transfer (BRET)^6^, and split enzyme complementation assays^7^. In the latter system, enzymes are dissected into two non-active components and fused to proteins that bind to an analyte on different epitopes. When these two components are brought into proximity by presence of the target, they reconstitute an active reporter enzyme.^8^ For instance, Stains et al. demonstrated the use of firefly luciferase complementation assay to detect HIV-1 gp120, human VEGF, and HER-2.^9^ In a similar approach, Mie et al. fused the two fragments of split *Renilla* luciferase to the B domain of protein A of *Staphylococcus aureus* and used it to detect *E. coli*.^10^ Split enzyme systems are advantageous due to their high sensitivity, simplicity, and speed.^8^

**Figure 1.**
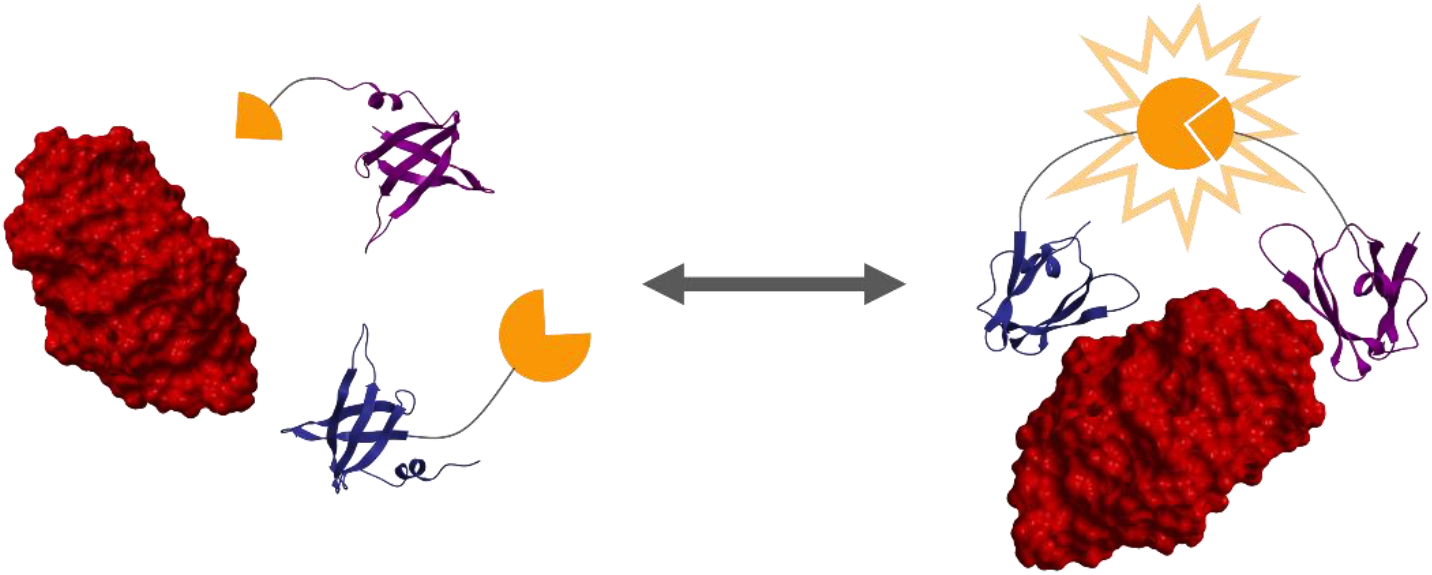
Split luciferase mix-and-read assay for the detection and quantification of a soluble target. Binders to the target molecule, lysozyme (PDB: 2CDS; red), in this case derived from the Sso7d scaffold protein (PDB 1SSO; blue and purple) are fused to N- or C-terminal fragments of split luciferase (orange) to create biosensor probes. Upon addition of the probes to a solution containing the target, probes bind to the target, and, because of the proximity created by these binding events, fragments of the split luciferase assemble to create an active luciferase enzyme. When substrate is added, a luminescent signal is produced corresponding to concentration of the target in solution.

In previous work, we have developed a mix-and-read split protein complementation assay for the detection of a model target, lysozyme^11^, based on the tripartite GFP system^12^. In that study, two binding proteins targeting epitopes on lysozyme were identified. These binding proteins, NTL1 and CTL1 were obtained by mutagenesis of the Sso7d protein from the hyperthermophilic archaeon *Sulfolobus solfataricus*. Sso7d is a small (7.4 kDa), highly stable and versatile scaffold that has been used to produce binding proteins to various targets, while retaining thermal and chemical stability.^13–15^ NTL1 and CTL1 bind to lysozyme on non-overlapping epitopes with equilibrium dissociation constants (*K*_*D*_s) ~ 1.3 μM and 250 nM, respectively. The binders were fused to two parts of a tripartite split GFP, which allowed for target-dependent fluorescence when the target and third GFP unit were present. The limit of detection (LoD) and limit of quantification (LoQ) of this assay were 235 nM and 460 nM, respectively. ^11^

In this work, we describe a framework to develop simple mix-and-read complementation assays for sensitive detection and quantification of soluble targets, by combining engineered protein binders with split luciferase. Due to their higher signal-to-noise ratio, split luciferase systems have advantages over assays using split fluorescent proteins.^16^ We also describe a simple equilibrium binding model for mix-and-read assays based on split luciferase. Specifically, the model provides an analytical framework to link binding affinities of the probes for the target, assay parameters such as the concentrations of probes used, and assay performance (limit of detection and concentration range over which the target can be quantified). Given preliminary detection data of a target molecule by the split luciferase assay, the model can predict the concentration of binding probes needed to accurately quantify the target in a particular concentration range. Due to our model and the ease of creating new Sso7d-based binders, our mix-and-read assay can be easily adapted to new targets or to detect different concentration ranges of an existing target.

## Methods

### Library screening and binder selection

A previously constructed library of Sso7d mutants was used for the isolation of binders (Snof3 and Snof10) for the model target protein Sso6904.^13^ The yeast surface display library was screened against recombinant Sso6904 as previously described, using one round of magnetic sorting and one round of fluorescence-activated cell sorting (FACS).^17^ Five unique high affinity binders were selected, and competitive binding experiments were performed to identify binders targeting non-overlapping epitopes on Sso6904. Briefly, yeast displaying binders 1, 5, 7, or 10 (Snof10) were labeled with Sso6904 and a 25-fold molar excess of soluble binder 3 (Snof3). Binding of Sso6904 to binders 1, 5, 7, or 10, in the presence of binder 3 was assessed with flow cytometry. Detailed protocols for yeast culture, library screening, and competitive binding experiments can be found in the Supplementary Methods.

### Estimation of K_D_

Apparent binding affinities of Snof3 and Snof10 to Sso6094 were estimated using yeast surface titrations and data was fit to a monovalent binding isotherm as previously described.^17^ Detailed protocols for yeast surface titrations and data fitting can be found in the Supplementary Methods.

### Expression and purification of Sso6904, Snof3, and split luciferase probes

The DNA sequences for Sso6904, Snof3, and all split luciferase probes were cloned into pET-28b(+) or pET-22b(+) plasmids and transformed into Rosetta *E. coli* cells for recombinant protein expression. These proteins were purified using ion exchange chromatography, immobilized metal affinity chromatography, or both. Detailed protocols for construction of the plasmid vectors containing these proteins and their expression and purification can be found in the Supplementary Methods.

### Split luciferase mix-and-read assay

Varying concentrations of target protein lysozyme (Sigma-Aldrich) or Sso6904 and constant, equal concentrations of two corresponding split luciferase probes (i.e., *P*_1*T*_ = *P*_2*T*_) were mixed in PBS containing 0.1% BSA in a total volume of 350 μL. The solutions were allowed to equilibrate at room temperature with rotation for the specified equilibration step time. 50 μL of the protein solution was transferred to a white, clear bottom 96-well plate in duplicate. Nano-Glo^®^ Luciferase Assay Reagent (Promega) was prepared according to the manufacturer’s protocol. 50 μL of the reagent was added to each well and the plate was incubated at room temperature with rotation for the specified detection incubation time. Luminescence was quantified on a microplate reader (Tecan) with a 1000 ms integration time. The experimental limits of blank, detection, and quantification were estimated as described in the Supplementary Methods.

### Estimation of apparent affinity of binding interaction between probes 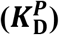

Apparent affinity 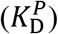 of binding interaction between the lysozyme detection probes, and between the Sso6904 detection probes were estimated by conducting split luciferase mix-and-read titration assays. Split luciferase assays were performed as described above with constant concentrations of one probe, varying amounts of the other probe, and no target protein. Data was fit to a modified binding isotherm.^17,18^ Detailed protocols for split luciferase mix-and-read titration assays and data fitting can be found in the Supplementary Methods.

### Model development and fitting

The split luciferase equilibrium binding model is represented mathematically by conservation equations, bimolecular equilibrium dissociation constants, and unimolecular equilibrium dissociation constants, all of which are shown in the Supplementary Methods. The assumption that linkers connecting the enzyme fragment and binding protein in each probe are in positions such that 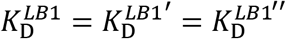, 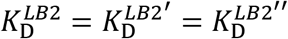, and 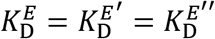 was made. We also assumed that the concentrations of *C*_*EO*_, *C*_*LB*1*O*_, and *C*_*LB*2*O*_ were negligible. Under these assumptions, the model equations were the following:

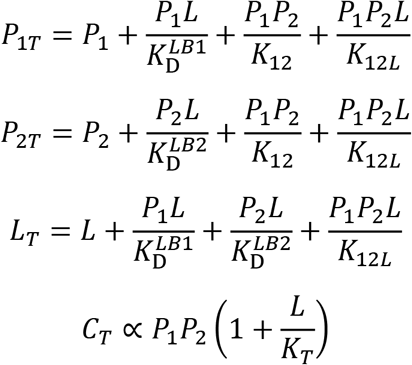

The first two equations are conservation equations for *P*_1*T*_ and *P*_2*T*_, the (known) total concentrations of probe 1 and probe 2 in the sample before mixing. The third equation is the conservation equation for *L*_*T*_, the combined concentration of all species containing the target molecule and equal to concentration of the target in the sample before mixing. In the final equation, *C*_*T*_ is equal to the concentration of all signal-producing complexes in a sample and is the model output. In the first three equations, terms that contain the product of *P*_1_ and *P*_2_ have the lumped parameter *K*_12_, and terms that contain the product of *P*_1_, *P*_2_, and *L* contain the lumped parameter *K*_12*L*_:

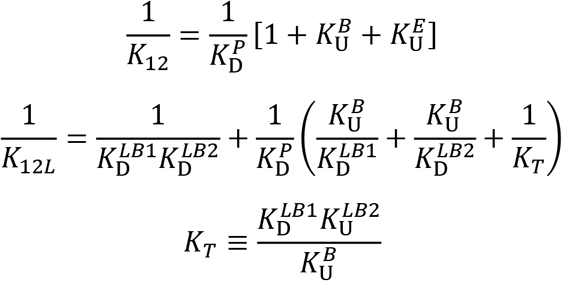

The model equations contain five parameters: 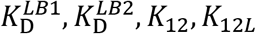, and *K*_*T*_. If 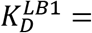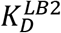 and if *P*_1*T*_ = *P*_2*T*_, then the model can be further simplified to three reduced model equations.

A two-stage fitting method was used to determine best fit values of all unknown parameters in the full and reduced model. A detailed description of the model creation and fitting the full and reduced model to experimental data can be found in the Supplementary Methods. Additionally, methods for averaging and normalization of the data to which the model was fit, estimation of experimental variance as a function of *C*_*T*_, and estimation of model-derived limits of blank, detection, and quantification can be found in the Supplementary Methods.

## Results and Discussion

### Development of a split luciferase assay for target detection

Split luciferase complementation systems have been created with firefly (*Photinus pyralis*) luciferase^19,20^, *Renilla* luciferase^21^, *Gaussia* luciferase^22^, and an engineered subunit of *Oplophorus gracilirostris* luciferase, termed NanoLuc^16,23^. NanoLuc is the engineered catalytic subunit of luciferase from the deep sea shrimp *Oplophorus gracilirostris* and is small in size (19 kDa), but possesses high thermal stability and 150-fold greater luminescence intensity than firefly and *Renilla* luciferase.^24^ Because of these qualities, we chose to use a split luciferase system based on NanoLuc luciferase over firefly and *Renilla* luciferase. The small size of NanoLuc makes it suitable for our complementation assay as the small NanoLuc fragments are less likely to disrupt the activity of their partner binding proteins by steric hindrance. Like NanoLuc, *Gaussia* luciferase has a low molecular weight and exhibits high emission intensity, however the luminescence of *Gaussia* luciferase decays rapidly under most conditions, and the *Gaussia* luciferase system can produce high autoluminescence background.^24^ Both factors would limit the sensitivity of a spilt luciferase assay based on *Gaussia* luciferase; therefore, we chose to use NanoLuc over *Gaussia* luciferase to develop our split luciferase assay.

We considered two versions of split NanoLuc for our complementation assay. Verhoef et al. developed one split NanoLuc system, referred to as sNLUC, by dividing NanoLuc after amino acid 52 creating an N-terminal fragment NLF1 (5.7 kDa) and a C-terminal fragment NLF2 (13.4 kDa).^16^ To construct the other system, Dixon et al. split NanoLuc after amino acid 156 and structurally optimized the resulting fragments to create N-terminal subunit LgBiT (18 kDa) and C-terminal subunit SmBiT (1.3 kDa).^23^ This system is referred to as NanoBiT.

To create the components of our split luciferase assay, we fused NTL1 and CTL1, the two Sso7d-derived lysozyme binders identified by Carlin et al., to the C-terminal and N-terminal fragments of the split NanoLuc systems, as illustrated in **Fig. 1**. This resulted in sNLUC-based lysozyme probes CTL1-NLF1 and NLF2-NTL1 and NanoBiT-based lysozyme probes CTL1-LgBiT and SmBiT-NTL1.

We used both split NanoLuc systems to develop split luciferase complementation assays and assess their ability to detect a range of concentrations of lysozyme (**Fig. 2**). 1 μM sNLUC probes or 1 μM NanoBiT probes were added to solutions containing lysozyme at different concentrations. The samples were allowed to equilibrate for one hour before luciferase substrate was added to each mixture and luminescence was quantified immediately. Luminescence was detected in all solutions where sNLUC or NanoBiT probes were present. The sNLUC system showed a linear response to lysozyme between 250 nM and 1500 nM (**Fig. 2A**). Initial assessment showed that luminescent signal was saturated with 1μM of the NanoBiT probes; therefore, their concentration was decreased to 100 nM. As seen in **Fig. 2B**, the linear range of the NanoBiT system with 100 nM probes was from 50 nM to 100 nM. These results demonstrate that lysozyme binders NTL1 and CTL1 can be used in a split luciferase mix-and-read assay and allow for the lysozyme-mediated reconstitution of both sets of split NanoLuc fragments.

**Figure 2.**
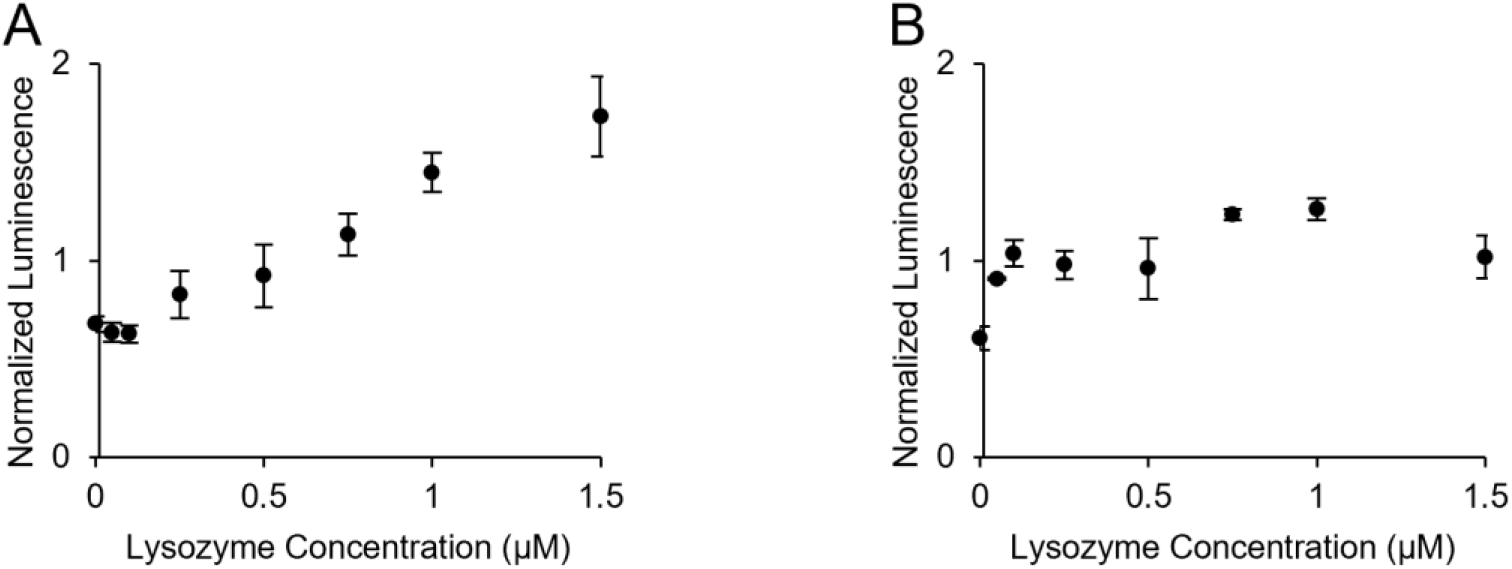
Split luciferase lysozyme detection assay with sNLUC and NanoBiT systems. Detection of lysozyme at different concentrations was measured with the mix-and-read split luciferase detection assay using either 1 μM sNLUC-based probes, CTL1-NLF1 and NLF2-NTL1 (A) or 100 nM NanoBiT-based probes, CTL1-LgBiT and SmBiT-NTL1 (B). Luminescence is normalized by the mean luminescent signal at all lysozyme concentrations for each repeat. Three independent replicates were conducted. Error bars indicate standard error.

The data in **Fig. 2** were used to quantify the limit of detection (LoD) and limit of quantification (LoQ) of the NanoBiT and sNLUC system. The LoD and LoQ of the NanoBiT assay (74 nM and 120 nM, respectively) were both over three times lower than the LoD and LoQ of the sNLUC assay (390 nM and 550 nM, respectively) and tripartite-GFP assay (235 nM and 460 nM, respectively).^11^ This suggests that lysozyme can be reliably detected at lower concentrations using the NanoBiT assay than the sNLUC or tripartite-GFP assays. This may be in part because of NanoBiT’s high signal to noise ratio compared to sNLUC, indicated by a lower limit of blank (LoB); the LoB of the NanoBiT and sNLUC assays were 34 nM and 240 nM.

It is important to note that the NanoBiT fragments were designed to minimize association with each other and have a binding affinity of 190 μM.^23^ This weak association is critical to ensure a low signal to noise ratio when the NanoBiT fragments are fused with binding proteins. Binding interaction between the split luciferase fragments combined with weak interactions between the two binder proteins may result in significant affinity of the two probes for each other. Indeed, the apparent affinity between the probes CTL1-LgBiT and SmBiT-NTL1 was 210 nM, despite the very low affinity of interaction between LgBiT and SmBiT (**Fig. S1**).

Because of its high sensitivity and low detection limit, we chose to further develop the NanoBiT-based split luciferase mix-and-read assay.

### Assay optimization

We hypothesized that we could further improve the split luciferase complementation assay to increase its sensitivity and dynamic range by tuning the probe concentrations. Specifically, we decreased the concentrations of probes CTL1-LgBiT and SmBiT-NTL1 to 1, 5, or 10 nM each. To minimize experimental variability arising from differences in incubation times, particularly during the detection step, we adjusted the equilibration and detection incubation time to four hours and one hour, respectively, to allow for equilibration of luminescent protein complexes in each sample. The equilibration step takes place after the probes are added but before the luciferase substrate is added to the sample with lysozyme. An additional incubation step, the detection incubation, is involved after the luciferase substrate is added prior to detection by chemiluminescence. Our results under these conditions are shown in **Fig. 3**.

**Figure 3.**
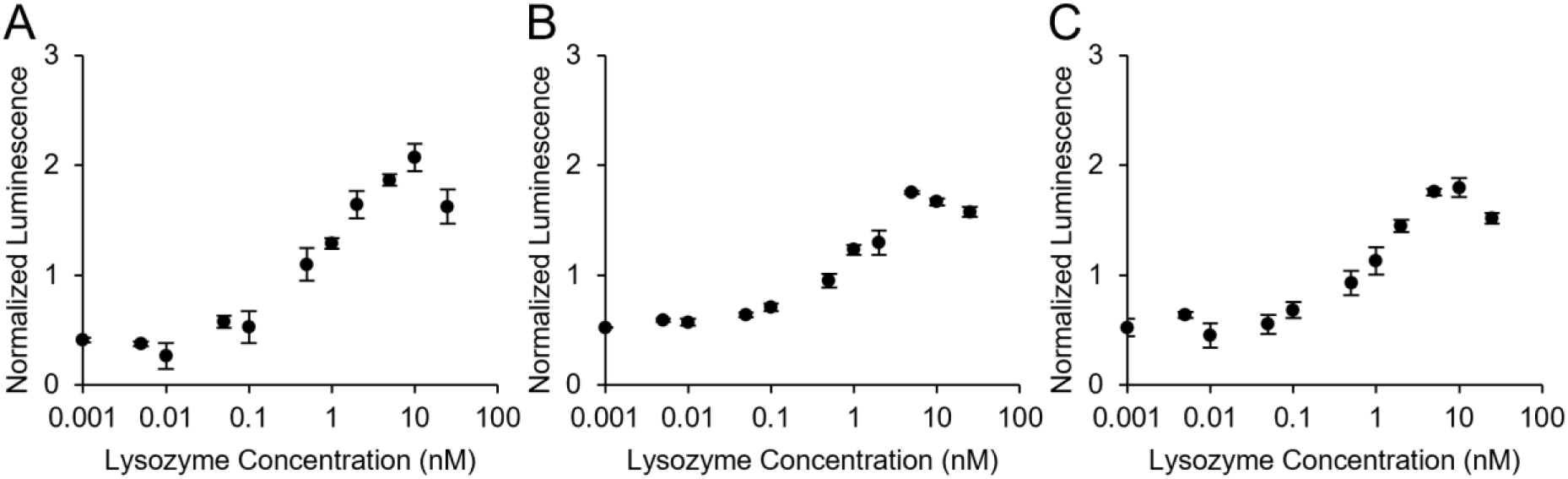
Optimized split luciferase lysozyme detection assay. Detection of lysozyme at different concentrations was measured with the mix-and-read split luciferase detection assay with 1 nM (A), 5 nM (B), or 10 nM (C) probes CTL1-LgBiT and SmBiT-NTL1. Luminescence is normalized by the mean luminescent signal at all lysozyme concentrations for each repeat. Four independent replicates were conducted at each probe concentration. Error bars indicate standard error.

With lower probe concentrations we observed an increase in assay sensitivity and reliable detection of picomolar concentrations of lysozyme. Lower concentrations of probes lead to less background noise, and consequently better assay sensitivity. Accordingly, the LoD and LoQ of the assay with 1 nM probes were lower than the LoD and LoQ with 5 nM or 10 nM probes, and were assessed as 29 pM and 37 pM, respectively. This corresponds to a greater than 2000-fold decrease in LoD and LoQ relative to the assays before optimization (**Fig. 2B** vs. **Fig. 3**). Reduction of background noise compared to the assay before optimization is illustrated by the low LoB (23 pM) with 1 nM probes—greater than 1000-fold less than the LoB resulting from the assay prior to optimization.

At lysozyme concentrations at or near the probe concentration, we observed a maximum in luminescent signal followed by a drop in signal (this maximum occurred at 10 nM with 1 nM probes). This trend was also seen, albeit less pronounced, with the NanoBiT-based lysozyme detection system before optimization. At analyte concentrations higher than the concentration of each probe, the probability that an analyte molecule will be bound by both probes decreases.^25^ Instead, only one of the two probes necessary for luciferase reconstitution may bind an analyte. Consequentially, luminescent output will underestimate the analyte concentration in each sample as seen in **Fig. 3**.

Unlike assay results prior to optimization, the system showed a linear response at low concentrations of lysozyme and a logarithmic response at higher concentrations. The linear and logarithmic ranges varied depending on the amount of probe used (**Fig. S2**), but with 1 nM probes were from 1 pM to 500 pM and from 500 pM to 10 nM, respectively. Accounting for the LoD and these ranges, the overall dynamic range^26^ of the assay is from 29 pM to 10 nM and covers 2.5 orders of magnitude, almost 20 times more than the dynamic range before optimization.

### Assay development for a new target

To assess the broader applicability, we evaluated the development of a split luciferase assay for a new target, where binding proteins targeting non-overlapping epitopes were not available. Specifically, we sought to assess if combinatorial library screening could be used to isolate binding proteins that bind non-overlapping epitopes on the target of interest, and if these binders could be incorporated into a split luciferase assay system for target detection.

Sso6904, an *S. solfataricus* protein, was chosen as a model target for our mix-and-read assay. First, we isolated a pair of novel binders targeting non-overlapping epitopes on Sso6904 from a library of 10^8^ Sso7d mutants by yeast surface display.^13^ To achieve this, we performed one round of magnetic sorting and one round of fluorescence-activated cell sorting (FACS) to screen the Sso7d library for Sso6904 binders. 18 Sso7d variants were randomly chosen from the post-FACS pool of binders and sequenced. Of these 18 mutants only 5 (binders 1, 3, 5, 7, and 10) were unique (**Fig. 4A**).

**Figure 4.**
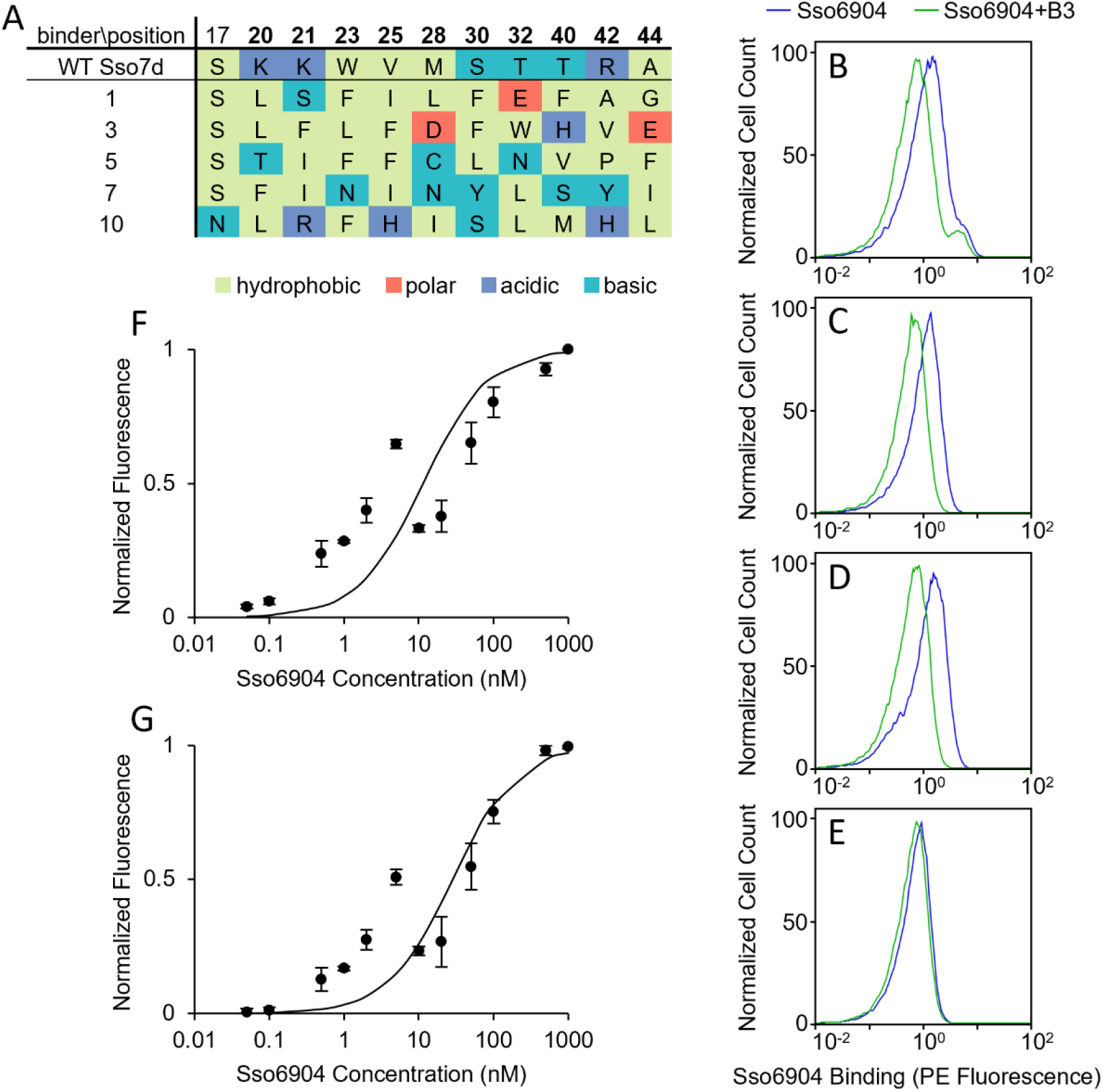
Identification of Sso7d-derived binders to non-overlapping Sso6904 epitopes. (A) Sequences of wild type Sso7d and unique binders to Sso6904 are shown. The ten positions mutated in the original Sso7d library are displayed in bold. (B-E) The results of the competitive binding experiments with soluble binder 3 and yeast surface displayed binder 1 (B), 5 (C), 7 (D), or 10 (E) are shown. Each flow cytometry plot depicts the normalized number of cells bound to Sso6904 via their surface displayed binder, measured by PE fluorescence, in the presence (green curve) or absence (blue curve) of excess binder 3. (F-G) Apparent *K*_*D*_s of Snof3 (F) and Snof10 (G) to Sso6904 were estimated using yeast surface titrations. Mean fluorescence was normalized by the maximum fluorescence of each repeat and *K*_*D*_ was calculated using a global non-linear least squares fit across three independent replicates for each binder. The *K*_*D*_ of Snof3 is 11 nM (68% confidence interval: 5.1 nM-24 nM) and the *K*_*D*_ of Snof10 is 28 nM (68% confidence interval: 17 nM-48 nM). Error bars correspond to standard error.

As shown in **Fig. 4A**, the composition of amino acids at the mutated residues in the binding site of binder 3 is unlike that of the other four mutants. None of the 10 mutated positions in binder 3 contain polar residues, while the other four mutants have an average of 2.5 polar residues and contain at least 1 polar amino acid in these 10 positions. Additionally, binder 3 contains the most residues mutated to an acidic amino acid (2), while only one other mutant has any acidic residues in these positions. These differences in the amino acid composition of the binding site of binder 3 led us to hypothesize that binder 3 may target Sso6904 at a distinct epitope from one of the other four binding proteins. To test our hypothesis, and to identify a pair of binding proteins that target distinct epitopes on Sso6904, we conducted competitive binding studies using binder 3 as a soluble competitor, as described below.

We incubated yeast displaying binder 1, 5, 7, or 10 with recombinant Sso604 in the presence or absence of a 25-fold excess of soluble binder 3 and assessed the binding of the surface displayed protein to Sso6904 with flow cytometry. **Figs. 4B-D** show that binder 3 outcompetes surface displayed binder 1, 5, or 7 for their binding site on Sso6904 resulting in a decrease in binding to Sso6904 in the presence of binder 3. Therefore, we concluded that binders 1, 3, 5, and 7 target overlapping epitopes on Sso6904. On the other hand, the addition of binder 3 to the sample containing yeast surface displayed binder 10 and Sso6904 had little to no effect on the ability of binder 10 to adhere to Sso6904 (**Fig. 4E**). Accordingly, we concluded that binders 3 and 10, hereafter referred to as Snof3 and Snof10, bind to Sso6904 on non-overlapping sites. We measured the binding affinities of the selected Sso6904 binding proteins using yeast surface titrations.^13^ The binding affinities of Snof3 and Snof10 for Sso6904 were 11 nM (68% confidence interval: 5.1 nM-24 nM) and 28 nM (68% confidence interval: 17 nM-48 nM), respectively (**Figs. 4F-G**).

### Detection of Sso6904

We used Snof3 and Snof10 to construct probes for the mix-and-read split luciferase assay to detect Sso6904, Snof3-LgBiT and SmBiT-Snof10. We performed split luciferase complementation assays, allowing the samples to equilibrate for four hours before adding the luciferase substrate and incubate for one hour before signal quantification, with 100 nM Snof3-LgBiT and 100 nM SmBiT-Snof10 (**Fig. 5**). Data from assays conducted in the same way with 10 nM of each probe are shown in the supplementary material (**Fig. S3**).

**Figure 5.**
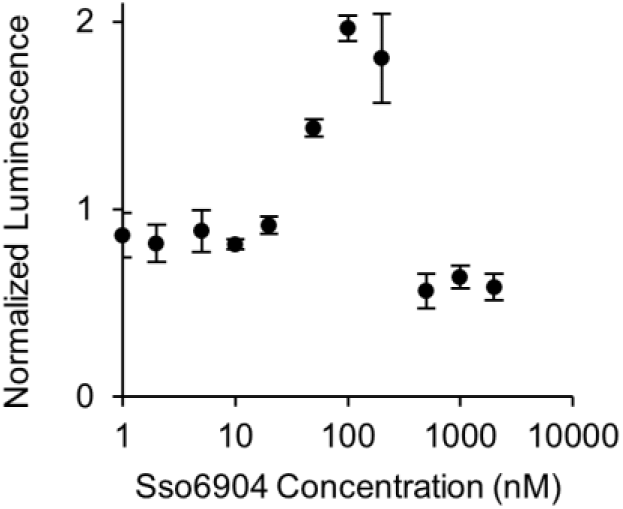
Split luciferase Sso6904 detection assay. Detection of Sso6904 at different concentrations was measured with the mix-and-read split luciferase detection assay using 100 nM Snof3-LgBiT and 100 nM SmBiT-Snof10. Luminescence is normalized by the mean luminescent signal at all Sso6904 concentrations for each repeat. Three independent replicates were conducted. Error bars indicate standard error.

Similar to assays for detection of lysozyme, luminescence was detected in all samples containing split luciferase probes. The linear range of Sso6904 detection with 100 nM probes was from 2 nM to 100 nM (**Fig. S2**). At Sso6904 concentrations higher than 100 nM, the concentration of probes used in the assay, we observed a drop in luminescent signal. This is consistent with prior results from the lysozyme detection system.

The results of these experiments illustrate that the Sso6904 probes allow for the concentration-dependent complementation of the split luciferase fragments and detection of Sso6904. Additionally, these results confirm that Snof3 and Snof10 bind to Sso6904 on non-overlapping epitopes. The LoB, LoD, and LoQ of the spilt luciferase Sso6904 detection assay are 21 nM, 40 nM, and 80 nM, respectively. The apparent affinity between Snof3-LgBiT and SmBiT-Snof10 was 150 nM (**Fig. S1**), indicating that the probes may bind to each other.

### A mathematical model for split luciferase detection of protein targets

Because the dynamic range of the split luciferase assay varies with probe concentration and the affinity of the binding component of each probe to the target molecule, we constructed an equilibrium binding model that relates these assay parameters and luminescent output to estimate the quantifiable range of the assay.^18^ The model can be used to estimate the concentrations of probes needed to quantify an analyte in a desired concentration range.

The equilibrium model describes all binding events occurring in our split luciferase detection system and is shown in **Fig. 6**. When constructing this model, we assumed that only the complexes shown in **Fig. 6** exist in our system. Unproductive complexes such as those containing more than one molecule of each probe or target do not occur. 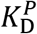, the apparent affinity between split luciferase probes, is defined by the relationship 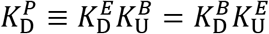. The equations used to model the system are shown in the Methods and Supplementary Methods.

**Figure 6.**
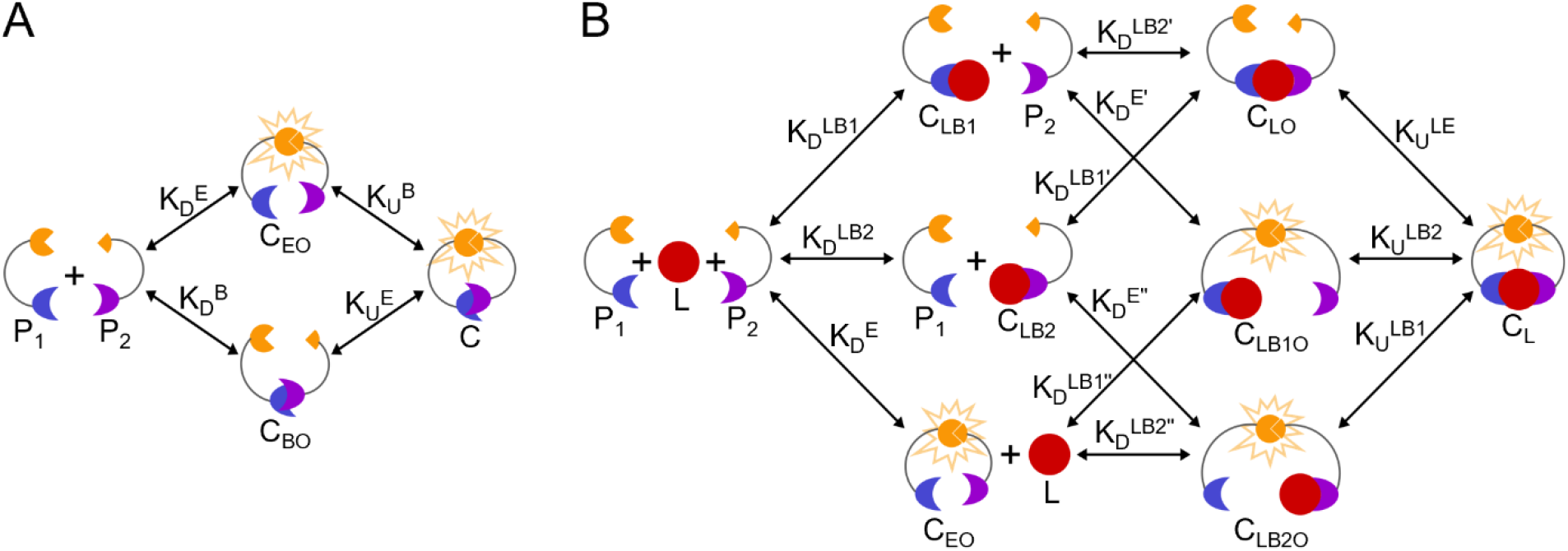
Proposed equilibrium model for the split luciferase assay. This model describes the split luciferase system when the target is not present (A) and when the target is present (B). Binder proteins (blue and purple); split luciferase fragments (orange); unbound target (*L*, red); unbound probe 1 (*P*_1_); unbound probe 2 (*P*_2_); luminescent, signal-producing complexes (*C*_*EO*_, *C*, *C*_*LB*1*O*_, *C*_*LB*2*O*_, and *C*_*L*_); non-luminescent complexes (*C*_*BO*_, *C*_*LB*1_, *C*_*LB*2_, and *C*_*LO*_); biomolecular dissociation constants; and unimolecular dissociation constants are shown.

The key equilibrium binding constants in this model are measurable and known for our lysozyme and Sso6904 detection systems. These include 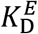 (190 μM^23^), 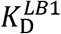 (1.3 μM^11^ for lysozyme detection and 28 nM for Sso6904 detection), 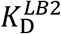 (250 nM^11^ for lysozyme detection and 11 nM for Sso6904 detection), and 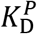 (210 nM for lysozyme detection and 150 nM for Sso6904 detection). Additionally, the concentration of probe 1 and probe 2 in the sample before mixing, *P*_1*T*_ and *P*_2*T*_, are known and are 1 nM, 5 nM, 10 nM, or 100 nM. *P*_1*T*_ and *P*_2*T*_ are also equal to the combined concentrations of all species containing probe 1 or probe 2. *L*_*T*_ is the combined concentrations of all species containing the target molecule and equal to concentration of the target in the sample before mixing. *C*_*T*_ is equal to the concentration of all signal-producing complexes in a sample.

To determine values of unknown parameters in the model, we fit our model to the results of the experimental split luciferase detection assays where *L*_*T*_, the initial concentration of target in each sample, and *C*_*T*_, a variable related to the assay’s luminescent output, were known. We fit our model to the results of the optimized lysozyme detection assay with 5 nM and 10 nM probes and to the results of the Sso6904 detection assay with 100 nM probes. Data from the detection of lysozyme with 1 nM probes was not included in the fit because the maximum luminescent signal from this assay did not occur at the concentration of lysozyme equal to the concentration of probes as seen in the other assays using NanoBiT-based probes.

The best model fit was determined by minimizing the errors between *C*_*T*_ calculated by the model and assay data (see **Supplementary Methods**). Model parameters were lumped and allowed to vary over several orders of magnitude that included ranges given by the confidence intervals of known, calculated parameters; estimates of unknown parameters; and estimates of the effective concentration during unimolecular binding events. Once values of these parameters were determined, the model equations were solved to produce an output luminescent signal (*C*_*T*_) given an initial target concentration (*L*_*T*_). The luminescence curves resulting from model equations solved with parameters fit to lysozyme detection data and 1 nM, 5 nM, and 10 nM input probe concentrations are shown in **Fig. 7A**. The curves generated from model equations solved with Sso6904 detection-derived parameters and 100 nM input probe concentrations is shown in **Fig. 7B**.

**Figure 7.**
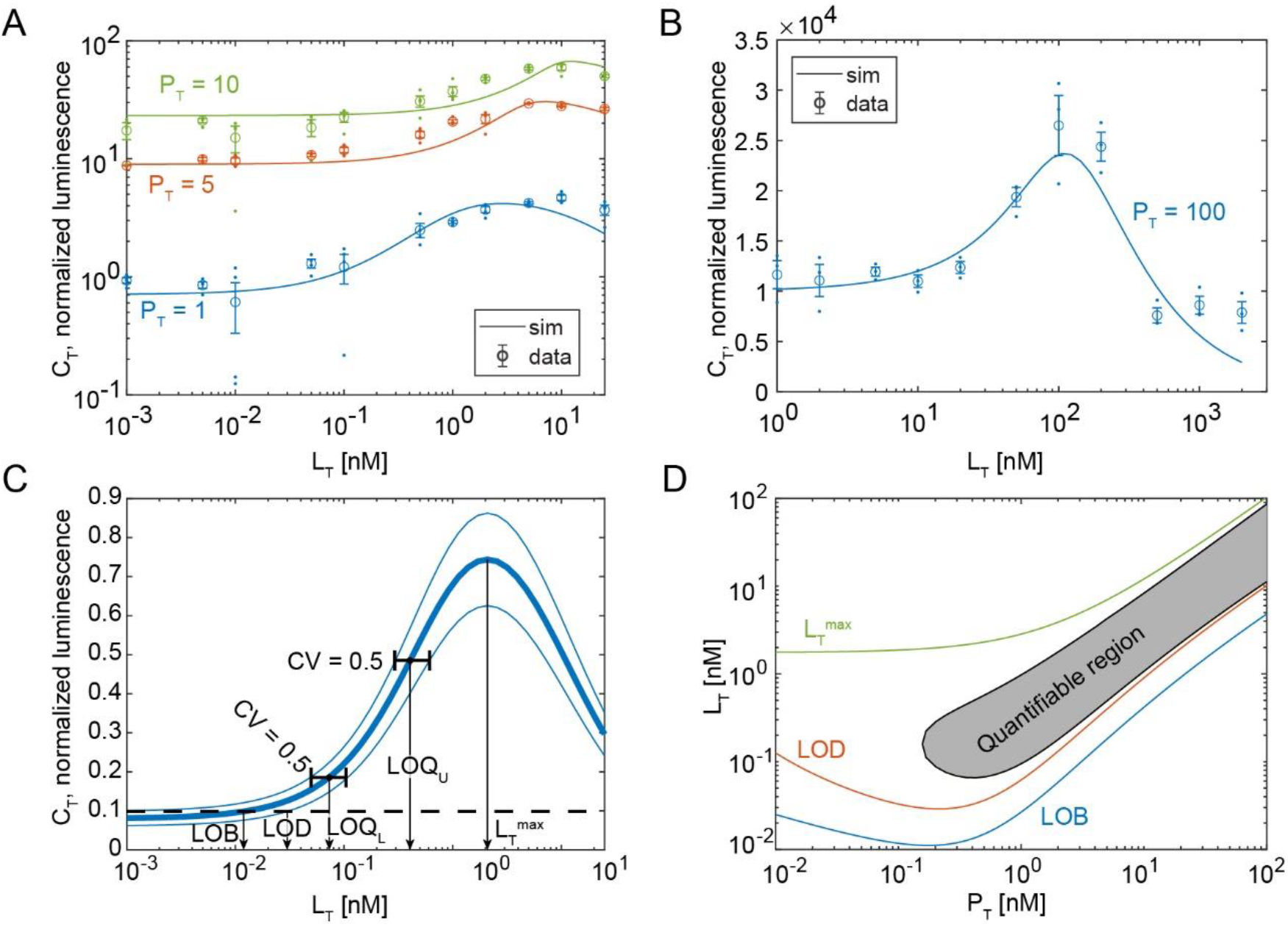
Output of equilibrium binding model. (A-B) Total luminescent output (*C*_*T*_) versus total target concentration (*L*_*T*_) curves given by the equilibrium model fit to split luciferase lysozyme detection assay data (A) or split luciferase Sso6904 detection assay data (B). Curves show model simulation, solid dots are three repeats of experimental data, and open circles are averages of experimental data. Luminescence was normalized by dividing data by maximum luminescent output, averaging these values at each target concentration over the three repeats to create an average curve, and applying a multiplier to each repeat to minimize the distance between it and the average curve. (C) Model derived *C*_*T*_ vs *L*_*T*_ curve and LoB, LoD, lower LoQ (*LoQ*_*L*_), upper LoQ (*LoQ*_*U*_), and *L*_*T*_ at the maximum luminescence (*L*_*T*_^*max*^) predictions for the lysozyme detection system at *P*_*T*_ = 0.3 nM. (D) Plot of how the LoB, LoD, quantifiable region, and *L*_*T*_^*max*^ vary with respect to probe concentration *P*_*T*_ for the lysozyme detection system.

The model we have constructed was able to fit data sets generated from the split luciferase mix-and-read assay detecting lysozyme and Sso6904 analytes. The model captures the linear-then-logarithmic trend observed in our lysozyme detection data and the linear response seen with Sso6904 detection. Overall, our model is capable of predicting output luminescence at low analyte concentrations, however in our experimental results, we observed a slight dip in signal at 10 pM lysozyme or 10 nM Sso6904 and this phenomenon is not seen in the curves produced by our model. Consistent with our assay results, the model shows a maximum luminescence occurring at or near the probe concentration in the system. The inflection point before this maximum luminescence value is not always accurately reflected by the model, as seen in the model curves for lysozyme detection with 5 nM and 10 nM probes in **Fig. 7A**.

The binding protein dissociation constants, 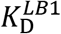 and 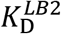, determined by the model to best fit the Sso6904 detection data were both 20 nM. This is almost exactly in between the experimentally determined values for 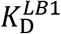 and 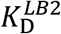, 28 nM and 11 nM, respectively. 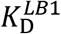 and 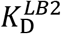 used by the model for the fitting of the lysozyme detection data were 1.8 nM, a value much lower than the independently measured binding affinities. There may be unaccounted for interactions occurring between the binders, enzyme fragments, or analyte molecule in the lysozyme detection system and not in the Sso6904 detection system making the model determined 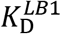 and 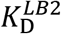 for the lysozyme system appear lower than the experimentally determined binding affinities and explaining this inconsistency.

From the error seen among biological repeats and technical error seen within biological repeats in the experimental data, we calculated biological variance and technical variance, respectively, as a function of *C*_*T*_. Biological and technical variance were combined to determine the total variance of the assay for a particular target as a function of *C*_*T*_. We used this total uncertainty to calculate the assay’s LoB, LoD, lower LoQ, and upper LoQ at any total probe concentration *P*_*T*_ where *P*_*T*_ = *P*_1*T*_ = *P*_2*T*_.^27,28^ Because the readout of the split luciferase assay has a maximum (*L*_*T*_^*max*^), there are two LoQs, a lower LoQ (*LoQ*_*L*_) and an upper LoQ (*LoQ*_*U*_). The model estimated LoB, LoD, lower and upper LoQ, and *L*_*T*_^*max*^ are shown for the lysozyme detection system at *P*_*T*_ = 0.3 nM in **Fig. 7C**. The area bounded by the lower and upper LoQs is the quantifiable region, or the range in which we are 95% confident that the measured *L*_*T*_ value is within two-fold of the true *L*_*T*_ and the range in which we can accurately quantify the target molecule. **Fig. 7D** illustrates how the LoB, LoD, quantifiable region, and *L*_*T*_^*max*^ vary with respect to probe concentration for the lysozyme detection system.

## Conclusion

The split luciferase mix-and-read assay and equilibrium binding model that we have developed enable the detection and quantification of various analytes in a sample at low picomolar concentrations. Our simple mix-and-read assay does not require multiple complicated steps and is broadly applicable to detect proteins or other molecules where protein binders can be developed. Using our optimized experimental setup for the detection of a model target we were able to quantify concentrations of the target spanning over 2.5 orders of magnitude starting at less than 50 pM of target protein. The multivalent nature of the interaction between the probes and target enables a highly sensitive assay even when using binding proteins with low to moderate affinities. This is also due in part to the low background noise produced in our system, demonstrated by a low LoB. However, gaining higher sensitivity required equilibration times to increase from one hour to four hours; therefore, there is a trade-off between speed and sensitivity.

Our lysozyme detection system was able to detect and quantify much lower concentrations of the target than our Sso6904 detection system despite the lower binding affinities of the Sso6904 binding proteins. One possible explanation for this is that NTL1 and CTL1 were screened together as a bivalent binder to lysozyme whereas Snof3 and Snof10 were selected as individual monovalent binders to Sso6904 without considering the proximity of their binding sites. Furthermore, the linkers connecting the two components of each probe were designed with the lysozyme detection system in mind. While the Sso6904 detection probes allow for the detection of Sso6904 in the nanomolar range, they may not bind to Sso6904 at positions or in orientations most optimal for luciferase reconstitution. On the other hand, the lysozyme detection probes were created to have the ideal size and binding sites to allow for reconstitution of the split luciferase fragments. Our results highlight the benefit of isolating a pair of binders together as a bivalent binder rather than separately.

Through the detection of lysozyme and Sso6904, we have shown that this assay can be quickly adapted for the quantification of multiple target molecules. Non-antibody binders can be easily isolated from combinatorial libraries with established protocols and plugged into our modular split luciferase system. We created an equilibrium binding model that defines a relationship between the luminescent output of the split luciferase assay, the concentration of biosensor probes used in the assay, and the binding affinities of the two binding proteins. Given data from a few detection assays (with varying probe and target concentrations), the model can predict the assay’s LoB, LoD, lower and upper LoQ, or quantifiable region of a target molecule, at any total probe concentration and can therefore predict the concentration of probes needed to quantify the target in a given concentration range. Thus, collectively our work provides the experimental and analytical framework to develop and optimize split luciferase-based assays for various protein targets.

