## Supplementary Information for "Experimental and analytical framework for “mix-and-read” assays based on split luciferase"

[2]Current address: School of Chemical and Biomolecular Engineering, Georgia Institute of Technology, Atlanta, GA 30332, USA

[3]Current address: Medical Scientist Training Program and Program in Genetics and Genomics, Baylor College of Medicine, Houston, TX 77030, USA

*Address correspondence to

### Contents

Supplementary Methods

References

Supplementary Tables

- Table S1
- Table S2

Supplementary Figures

- Figure S1
- Figure S2
- Figure S3

### Supplementary Methods

Yeast culture

Sso7d mutants were displayed as C-terminal fusions to the cell wall protein Aga2 using the pCTCON vector which contains a tryptophan selectable marker. All yeast were cultured in tryptophan deficient SDCAA media (20 g/L dextrose, 6.7 g/L yeast nitrogen base, 5 g/L casamino acids, 5.4 g/L $Na_2HPO_4$, 8.6 g/L $NaH_2PO_4 \cdot H_2O$) at 30°C with shaking at 250 rpm. To induce cell surface protein expression, yeast cells were transferred to tryptophan deficient SGCAA media (20 g/L galactose, 2 g/L dextrose, 6.7 g/L yeast nitrogen base, 5 g/L casamino acids, 5.4 g/L $Na_2HPO_4$, 8.6 g/L $NaH_2PO_4 \cdot H_2O$) at an $OD_{600}$ of 1 and cultured overnight at 20°C with shaking at 250 rpm.

Screening a combinatorial Sso7d library against Sso6904

The Sso7d library was screened against recombinant Sso6904 as previously described using yeast surface display and one round of magnetic sorting and one round of fluorescence-activated cell sorting (FACS).[1] The magnetic sort was performed against magnetic Dynabeads® Biotin Binder beads coated with biotinylated Sso6904. Sso6904 was biotinylated with EZ-Link™ Sulfo-NHS-LC-Biotin following the manufacturer's protocol and using at 10-fold molar excess of biotin. Excess reagent was removed by dialysis in tris buffer (50 mM Tris-HCl, 300 mM NaCl pH 7.4). Sso6904 coated beads were prepared by washing $10^7$ beads twice with PBS containing 0.1% BSA (PBS-BSA) using a magnet and incubating washed beads with 20 μg biotinylated Sso6904 in 500 μL PBS-BSA overnight at 4°C with rotation. Following overnight incubation, the beads were washed with PBS-BSA and resuspended in 100 μL PBS-BSA.

$10^9$ induced library yeast cells were washed twice and resuspended in 4 mL PBS-BSA. Two negative selections against Dynabeads® Biotin Binder beads and one positive selection against Sso6904-coated beads were performed. For the first negative selection, $10^7$ washed beads were added to the library cells and incubated for 1 hour at 4°C with rotation. Unbound yeast cells were collected, and the negative selection was repeated with $10^7$ new beads. After the second negative selection, yeast cells were incubated with $10^7$ Sso6904-coated beads for 1 hour at 4°C with rotation. Unbound cells were discarded, and the beads were washed 5 times with PBS-BSA. The beads were transferred to 5 mL of SDCAA media and yeast bound to the beads were cultured for 36-48 hours.

Yeast cells from the magnetic sort were induced and labeled for FACS. $10^7$ induced cells were washed with PBS-BSA and simultaneously labeled with 100 nM biotinylated Sso6904 and a 1:100 dilution of chicken anti-c-myc antibody in 100 μL PBS-BSA for 1 hour on ice. Cells were washed with PBS-BSA, and secondary labeling was performed with a 1:250 dilution of streptavidin R-phycoerythrin conjugate and Alexa Fluor 488 goat anti-chicken antibody in 100 μL PBS-BSA for 10 minutes on ice. Cells were washed with PBS-BSA and sorted on a MoFlo Cell Sorter (Beckman Coulter). Yeast expressing high affinity binders to Sso6904 were collected and expanded in 5 ml SDCAA media for 36-48 hours.

The resulting cells were plated on SDCAA plates (20 g/L dextrose, 6.7 g/L yeast nitrogen base, 5 g/L casamino acids, 5.4 g/L $Na_2HPO_4$, 8.6 g/L $NaH_2PO_4 \cdot H_2O$, 182 g/L sorbitol, 12 g/L agar) and allowed to grow for 36-48 hours at 30°C. 18 yeast colonies were selected and used to inoculate 5 mL SDCAA cultures. After 24-48 hours, plasmid DNA was extracted from each yeast culture using a Zymoprep™ Yeast Plasmid Miniprep II kit. The plasmids were each transformed into electrocompetent NovaBlue *E. coli* cells and the *E. coli* was grown overnight in a 5 mL LB (10 g/L tryptone, 5 g/L yeast extract, 10 g/L NaCl) culture at 37°C with shaking at 250 rpm. Plasmid DNA was extracted and sequenced.

Competitive binding experiments

Yeast cells displaying binders 1, 5, 7, or 10 (Snof10) on their surface were cultured and induced overnight in SGCAA media. $2 \times 10^6$ of each cell type were washed with PBS-BSA. The yeast were labeled with 2 nM biotinylated Sso6904 in the presence or absence of 50 nM binder 3 (Snof3) in 50 µL PBS-BSA for 20 minutes with rotation at room temperature. Cells were washed with PBS-BSA and labeled with a 1:250 dilution of streptavidin R-phycoerythrin conjugate in 50 µL PBS-BSA for 10 minutes on ice. Cells were washed with PBS-BSA and Sso6904 binding was assessed with flow cytometry

Estimation of Snof3 and Snof10 $K_D$s

Apparent binding affinities of Snof3 and Snof10 to Sso6094 were estimated using yeast surface titrations and data was fit to a monovalent binding isotherm as described.[1] $10^6$ yeast cells expressing Snof3 or Snof10 were labeled with varying concentrations of biotinylated Sso6904 and subsequently with a 1:250 dilution of streptavidin R-phycoerythrin conjugate. Primary labeling with biotinylated Sso6904 was performed in volumes to ensure ~10-fold or greater excess of Sso6904 target molecules over surface displayed binders. The fraction of cell surface mutants bound to Sso6904 was assessed with flow cytometry. Yeast surface titrations and flow cytometric analysis were performed 3 times for each binder. Background-subtracted mean PE fluorescence was normalized by the background-subtracted maximum mean PE fluorescence value for each of the three repeats to obtain the normalized fluorescence, $F$. Normalized fluorescence was related to initial target concentration, $[L]_0$, using the following relationship

$$F = \frac{[L]_0}{K_D + [L]_0}$$

Experimentally obtained data were fit to this equation using global non-linear least squares regression and a single $K_D$ value for each binder was determined as a fitted parameter across all three repeats.

Construction of plasmids containing Sso6904, Snof3, and split luciferase probes

All gene fragment and primer sequences used for cloning can be found in Supplementary Tables 1 and 2, respectively. Gene Fragment 1, a gene fragment containing the Sso6904 sequence, and the plasmid pET-22b(+) were digested with NdeI and XhoI restriction enzymes and Sso6904 was ligated into pET-22b(+). The Snof3 sequence was amplified by PCR with primers Pf1 and Pr1 from the pCTCON vector containing Snof3 extracted from yeast. The Snof3 PCR product and pET-22b(+) were digested with NdeI and XhoI and Snof3 was ligated into pET-22b(+).

The vector pCTCON containing lysozyme binders NTL1 and CTL1 connected with a linker was used as a backbone for the initial cloning steps to create plasmids containing our split luciferase probes. The construction of this vector is described elsewhere.[2] *Gaussia* luciferase surrounded by N- and C-terminal linkers (Gene Fragment 2) was amplified by PCR with primers Pf2 and Pr2 to introduce SmaI and AvrII restriction sites into the gene. pCTCON containing NTL1 and CTL1 was digested with SmaI and AvrII and ligated with digested *Gaussia* luciferase PCR product to insert the *Gaussia* luciferase gene between NTL1 and CTL1. The *Gaussia* luciferase gene was then replaced with the NanoLuc gene. NanoLuc was amplified from Gene Fragment 3 with primers Pf3 and Pr3. pCTCON containing the lysozyme binders and *Gaussia* luciferase and the NanoLuc gene were both digested with NsiI and SpeI and then ligated together. This replaced *Gaussia* luciferase between NTL1 and CTL1 with NanoLuc.

The probe CTL1-NLF1 was generated by amplifying the CTL1 sequence and NanoLuc residues 1-52 with primers Pf4 and Pr4 from the pCTCON vector containing the lysozyme binders and NanoLuc. This fragment was then cloned into pET-28b(+) between NcoI and XhoI restriction sites. Similarly, NLF2-NTL1 was generated by amplifying NanoLuc residues 53-173 and NTL1 with primers Pf5 and Pr5 from the same vector and cloning this fragment into pET-22b(+) between NdeI and XhoI restriction sites.

The probe containing LgBiT was initially created with a different Sso7d mutant that does not target lysozyme. This construct was amplified from Gene Fragment 4 with Pf6 and Pr6 and cloned into pET-22b(+) between XbaI and XhoI restriction sites. The other Sso7d mutant was then replaced with CTL1. CTL1 was amplified from pCTCON containing the lysozyme binders with Pf7 and Pr7. The other Sso7d mutant was removed at NheI and BamHI restriction sites and CTL1 was cloned in between these sites. SmBiT-NTL1 was created by amplification of Gene Fragment 5 with primers Pf8 and Pr5 and cloned into pET-22b(+) between NdeI and XhoI restriction sites.

Snof3-LgBiT was generated by amplification of Gene Fragment 6 with Pf9 and Pr9 and cloned into pET-22b(+) between NdeI and XhoI restriction sites. To create SmBiT-Snof10, the sequence of Snof10 was amplified with Pf10 and Pr10 from the pCTCON vector containing Snof10 extracted from yeast. The Snof10 sequence and pET-22b(+) containing SmBiT-NTL1 were both digested with AvrII and XhoI and ligated together. This replaced the lysozyme binder NTL1 with the Sso6904 binder Snof10.

All PCR reactions were performed in 50 μL reactions with high-fidelity Phusion polymerase according to the manufacturer’s protocol. Restriction enzymes were purchased from New England Biolabs and ligations were performed with T4 DNA ligase. Plasmids were transformed into competent NovaBlue *E. coli* cells. Gene fragments were purchased from Addgene or Integrated DNA Technologies and primers were purchased from Integrated DNA Technologies or Eton Biosciences.

Expression and purification of Sso6904, Snof3, and split luciferase probes

Sso6904 in pET-22b(+), Snof3 in pET-22b(+), and the split luciferase probes, in either pET-22b(+) or pET-28b(+), were transformed into Rosetta *E. coli* cells for protein expressions. Cells were grown overnight in a 5 mL culture which was then used to inoculate a 1L 2XYT culture (16 g/L tryptone, 10 g/L yeast extract, 5 g/L NaCl). This culture was incubated at 37°C with shaking at 250 rpm until it reached an $OD_{600}$ between 0.6 and 0.8. Protein production was induced with 0.5 mM isopropyl β-D-1-thiogalactopyranoside (IPTG) and expression was carried out overnight at 20°C.

Proteins were purified using cation exchange (CEX) chromatography, anion exchange (AEX) chromatography, immobilized metal affinity chromatography (IMAC), or a combination of two of these methods using a BioLogic LP FPLC system (Bio-Rad). Cells were lysed in either 35 mL IMAC Buffer A (50 mM Tris-HCl, 300 mM NaCl, pH 8) or 35 mL IEX Buffer A (50 mM Tris-HCl, 50 mM NaCl, pH 8), and the soluble fraction of the cell lysate was loaded onto the appropriate 5 mL chromatography column. A Mini Profinity IMAC column (Bio-Rad), Mini Macro-Prep High S CEX column (Bio-Rad), and Mini Macro-Prep High Q AEX column (Bio-Rad) were used. The column was washed with 40 mL of IMAC Buffer A for IMAC or 40 mL IEX Buffer A for CEX or AEX chromatography. Protein was eluted with a 0-100% linear gradient of IMAC Buffer B (50 mM Tris-HCl, 300 mM NaCl, 500 mM imidazole, pH 8) or a 0-50% linear gradient of IEX Buffer B (50 mM Tris-HCl, 2 M NaCl, pH 8).

Fractions were analyzed by sodium dodecyl sulphate-polyacrylamide gel electrophoresis (SDS-PAGE) and fractions containing the pure protein of interest were combined and dialyzed into PBS pH 7.4 or tris buffer (25 mM Tris-HCl, 150 mM NaCl pH 7.4) as indicated. If the fractions did not contain the pure protein of interest after the first chromatography step, a second chromatography step was performed as described. Fractions containing the pure protein of interest, determined by SDS-PAGE, were combined and dialyzed into the appropriate buffer.

Sso6904 was purified by IMAC and dialyzed into PBS. Snof3 was purified first by CEX chromatography and fractions containing Snof3 were subsequently combined and purified by IMAC. Snof3 was dialyzed into PBS.

Split luciferase probes CTL1-NLT1 and CTL1-LgBiT were purified using the same protocol. These probes were purified by IMAC. If fractions did not contain pure probes after IMAC, they were combined and dialyzed into IEX Buffer A. An AEX chromatography step was performed, and the probes were dialyzed into tris buffer. NLF2-NTL1 and SmBiT-NLT1 were purified using the same method as their corresponding CTL1-fusions probes, except CEX was performed instead of AEX for the second chromatography step.

Snof3-LgBiT was purified first by AEX chromatography with a 0-100% IEX Buffer B gradient instead of a 0-50% gradient, and fractions containing Snof3-LgBiT were then combined and purified by IMAC. Snof3-LgBiT was dialyzed into tris buffer. SmBiT-Snof10 was purified by IMAC and dialyzed into tris buffer.

The concentrations of all proteins were quantified by bicinchoninic acid (BCA) assay.

Estimation of experimental limit of blank, detection, and quantification

Limit of blank (LoB), limit of detection (LoD), and limit of quantification (LoQ) were estimated using the following equations.[2,3]

$$LoB = M_B + 1.645(SD_B)$$

$$LoD = LoB + 1.645(SD_B)$$

$$LoQ = M_B + 5(SD_B)$$

…where $M_B$ is the mean luminescence of a sample without analyte, and $SD_B$ is the standard deviation of a sample without analyte.

Estimation of apparent affinity between probes, $K_{\mathrm{D}}^{P}$

Apparent affinity ($K_{\mathrm{D}}^{P}$) between the lysozyme detection probes CTL1-LgBiT and SmBiT-NTL1 and between the Sso6904 detection probes Snof3-LgBiT and SmBiT-Snof10 were estimated by conducting split luciferase mix-and-read titration assays. Split luciferase assays were performed as described with a constant concentration (25 nM) of one probe (SmBiT-NTL1 or Snof3-LgBiT), varying amounts of the other probe (CTL1-LgBiT or SmBiT-Snof10), and no target protein. Equilibration step and detection incubation times were 4 hours and 1 hour, respectively. Experiments were performed in triplicate. $K_{\mathrm{D}}^{P}$ was estimated from this data similarly to how $K_D$ was estimated for Snof3 and Snof10 except a correction in the isotherm equation was made because the one probe was not added in great excess of the other.[1,4] Background-subtracted mean luminescence was normalized by the background-subtracted maximum mean luminescence value for each of the three repeats to obtain the normalized luminescence, $y$. Normalized luminescence was related to the concentration of the probes using the following relationship

$$y = \frac{[L]_0 - y[P]_0}{K_{\mathrm{D}}^{P} + [L]_0 - y[P]_0}$$

…where $[L]_0$ is the concentration of the varying probe and $[P]_0$ is the concentration of the constant probe. Solving this equation for $y$ gives

$$y = \frac{(K_{\mathrm{D}}^{P} + [L]_0 + [P]_0) - \sqrt{(K_{\mathrm{D}}^{P} + [L]_0 + [P]_0)^2 - 4[P]_0[L]_0}}{2[P]_0}$$

Experimentally obtained data were fit to this equation using global non-linear least squares regression and a single $K_{\mathrm{D}}^{P}$ value for each set of probes was determined as a fitted parameter across all three repeats.

Model development

The split luciferase equilibrium binding model is represented mathematically by conservation equations,

$$P_{1T} = [P_1] + [C_{EO}] + [C_{BO}] + [C] + [C_{LB1}] + [C_{LO}] + [C_{LB1O}] + [C_{LB2O}] + [C_L]$$
$$P_{2T} = [P_2] + [C_{EO}] + [C_{BO}] + [C] + [C_{LB2}] + [C_{LO}] + [C_{LB1O}] + [C_{LB2O}] + [C_L]$$
$$L_T = [L] + [C_{LB1}] + [C_{LB2}] + [C_{LO}] + [C_{LB1O}] + [C_{LB2O}] + [C_L]$$
$$C_T = [C_{EO}] + [C] + [C_{LB1O}] + [C_{LB2O}] + [C_L]$$

bimolecular equilibrium dissociation constants,

$$K_{\mathrm{D}}^{E} = \frac{[P_1][P_2]}{[C_{EO}]}$$
$$K_{\mathrm{D}}^{B} = \frac{[P_1][P_2]}{[C_{BO}]}$$
$$K_{\mathrm{D}}^{LB1} = \frac{[P_1][L]}{[C_{LB1}]}$$
$$K_{\mathrm{D}}^{LB2} = \frac{[P_2][L]}{[C_{LB2}]}$$
$$K_{\mathrm{D}}^{LB2'} = \frac{[C_{LB1}][P_2]}{[C_{LO}]}$$
$$K_{\mathrm{D}}^{E'} = \frac{[C_{LB1}][P_2]}{[C_{LB1O}]}$$
$$K_{\mathrm{D}}^{LB1'} = \frac{[P_1][C_{LB2}]}{[C_{LO}]}$$
$$K_{\mathrm{D}}^{E''} = \frac{[P_1][C_{LB2}]}{[C_{LB2O}]}$$
$$K_{\mathrm{D}}^{LB1''} = \frac{[C_{EO}][L]}{[C_{LB1O}]}$$
$$K_{\mathrm{D}}^{LB2''} = \frac{[C_{EO}][L]}{[C_{LB2O}]}$$

and unimolecular equilibrium dissociation constants,

$$K_{\mathrm{U}}^{B} = \frac{[C_{EO}]}{[C]}$$
$$K_{\mathrm{U}}^{E} = \frac{[C_{BO}]}{[C]}$$
$$K_{\mathrm{U}}^{LE} = \frac{[C_{LO}]}{[C_L]}$$
$$K_{\mathrm{U}}^{LB2} = \frac{[C_{LB1O}]}{[C_L]}$$
$$K_{\mathrm{U}}^{LB1} = \frac{[C_{LB2O}]}{[C_L]}$$

The assumption that linkers connecting the enzyme fragment and binding protein in each probe are in positions such that $K_{\mathrm{D}}^{LB1} = K_{\mathrm{D}}^{LB1'} = K_{\mathrm{D}}^{LB1''}$, $K_{\mathrm{D}}^{LB2} = K_{\mathrm{D}}^{LB2'} = K_{\mathrm{D}}^{LB2''}$, and $K_{\mathrm{D}}^{E} = K_{\mathrm{D}}^{E'} = K_{\mathrm{D}}^{E''}$ was made. We also assumed that the concentrations of $C_{EO}$, $C_{LB1O}$, and $C_{LB2O}$ were negligible.

Rearranging and combining the equations above gives new model equations:

$$P_{1T} = P_1 + \frac{P_1 L}{K_{\mathrm{D}}^{LB1}} + \frac{P_1 P_2}{K_{\mathrm{D}}^{E}} + \frac{K_{\mathrm{U}}^{E} P_1 P_2}{K_{\mathrm{D}}^{E} K_{\mathrm{U}}^{B}} + \frac{P_1 P_2}{K_{\mathrm{D}}^{E} K_{\mathrm{U}}^{B}} + \frac{P_1 P_2 L}{K_{\mathrm{D}}^{LB1} K_{\mathrm{D}}^{LB2}} + \frac{P_1 P_2 L}{K_{\mathrm{D}}^{E} K_{\mathrm{D}}^{LB1}} + \frac{P_1 P_2 L}{K_{\mathrm{D}}^{E} K_{\mathrm{D}}^{LB2}} + \frac{P_1 P_2 L}{K_{\mathrm{D}}^{LB1} K_{\mathrm{U}}^{LB2} K_{\mathrm{D}}^{E}}$$

$$P_{2T} = P_2 + \frac{P_2 L}{K_{\mathrm{D}}^{LB2}} + \frac{P_1 P_2}{K_{\mathrm{D}}^{E}} + \frac{K_{\mathrm{U}}^{E} P_1 P_2}{K_{\mathrm{D}}^{E} K_{\mathrm{U}}^{B}} + \frac{P_1 P_2}{K_{\mathrm{D}}^{E} K_{\mathrm{U}}^{B}} + \frac{P_1 P_2 L}{K_{\mathrm{D}}^{LB1} K_{\mathrm{D}}^{LB2}} + \frac{P_1 P_2 L}{K_{\mathrm{D}}^{E} K_{\mathrm{D}}^{LB1}} + \frac{P_1 P_2 L}{K_{\mathrm{D}}^{E} K_{\mathrm{D}}^{LB2}} + \frac{P_1 P_2 L}{K_{\mathrm{D}}^{LB1} K_{\mathrm{U}}^{LB2} K_{\mathrm{D}}^{E}}$$

$$L_T = L + \frac{P_1 L}{K_{\mathrm{D}}^{LB1}} + \frac{P_2 L}{K_{\mathrm{D}}^{LB2}} + \frac{P_1 P_2 L}{K_{\mathrm{D}}^{LB1} K_{\mathrm{D}}^{LB2}} + \frac{P_1 P_2 L}{K_{\mathrm{D}}^{E} K_{\mathrm{D}}^{LB1}} + \frac{P_1 P_2 L}{K_{\mathrm{D}}^{E} K_{\mathrm{D}}^{LB2}} + \frac{P_1 P_2 L}{K_{\mathrm{D}}^{LB1} K_{\mathrm{U}}^{LB2} K_{\mathrm{D}}^{E}}$$

$$C_T = C + C_L = \frac{1}{K_{\mathrm{D}}^{E}} \left( \frac{P_1 P_2}{K_{\mathrm{U}}^{B}} + \frac{P_1 P_2 L}{K_{\mathrm{D}}^{LB1} K_{\mathrm{U}}^{LB2}} \right)$$

Note that, for the sake of brevity, the square brackets that denote concentration have been omitted. The above equations can be grouped by like terms:

$$P_{1T} = P_1 + \frac{P_1 L}{K_{\mathrm{D}}^{LB1}} + \frac{P_1 P_2}{K_{12}} + \frac{P_1 P_2 L}{K_{12L}}$$

$$P_{2T} = P_2 + \frac{P_2 L}{K_{\mathrm{D}}^{LB2}} + \frac{P_1 P_2}{K_{12}} + \frac{P_1 P_2 L}{K_{12L}}$$

$$L_T = L + \frac{P_1 L}{K_{\mathrm{D}}^{LB1}} + \frac{P_2 L}{K_{\mathrm{D}}^{LB2}} + \frac{P_1 P_2 L}{K_{12L}}$$

$$C_T \propto P_1 P_2 \left( 1 + \frac{L}{K_T} \right)$$

…where:

$$\frac{1}{K_{12}} = \frac{1}{K_{\mathrm{D}}^{E}} + \frac{K_{\mathrm{U}}^{E}}{K_{\mathrm{D}}^{E}K_{\mathrm{U}}^{B}} + \frac{1}{K_{\mathrm{D}}^{E}K_{\mathrm{U}}^{B}} = \frac{1}{K_{\mathrm{D}}^{E}K_{\mathrm{U}}^{B}}\left[1 + K_{\mathrm{U}}^{B} + K_{\mathrm{U}}^{E}\right] = \frac{1}{K_{\mathrm{D}}^{P}}\left[1 + K_{\mathrm{U}}^{B} + K_{\mathrm{U}}^{E}\right]$$

$$K_{\mathrm{D}}^{P} \equiv K_{\mathrm{D}}^{E}K_{\mathrm{U}}^{B} = K_{\mathrm{D}}^{B}K_{\mathrm{U}}^{E}$$

$$\frac{1}{K_{12L}} = \frac{1}{K_{\mathrm{D}}^{LB1}K_{\mathrm{D}}^{LB2}} + \frac{1}{K_{\mathrm{D}}^{E}K_{\mathrm{D}}^{LB1}} + \frac{1}{K_{\mathrm{D}}^{E}K_{\mathrm{D}}^{LB2}} + \frac{1}{K_{\mathrm{U}}^{LB2}K_{\mathrm{D}}^{E}K_{\mathrm{D}}^{LB1}}$$

$$= \frac{1}{K_{\mathrm{D}}^{LB1}K_{\mathrm{D}}^{LB2}} + \frac{1}{K_{\mathrm{D}}^{E}K_{\mathrm{U}}^{B}}\left(\frac{K_{\mathrm{U}}^{B}}{K_{\mathrm{D}}^{LB1}} + \frac{K_{\mathrm{U}}^{B}}{K_{\mathrm{D}}^{LB2}} + \frac{1}{K_{T}}\right)$$

$$K_{T} \equiv \frac{K_{\mathrm{D}}^{LB1}K_{\mathrm{U}}^{LB2}}{K_{\mathrm{U}}^{B}}$$

Given these equations, we have the following 5 parameters: $K_{\mathrm{D}}^{LB1}, K_{\mathrm{D}}^{LB2}, K_{12}, K_{12L}$, and $K_{T}$.

If $K_{D}^{LB1} = K_{D}^{LB2}$ and if $P_{1T} = P_{2T}$, then the symmetry of the experimental set up requires that $[P_1] = [P_2]$. Thus, the model equations reduce to:

$$P_{T} = P + \frac{PL}{K_{D}^{LB1}} + \frac{P^2}{K_{12}} + \frac{P^2 L}{K_{12L}}$$

$$L_{T} = L + 2\frac{PL}{K_{D}^{LB1}} + \frac{P^2 L}{K_{12L}}$$

$$C_{T} = \alpha P^2\left(1 + \frac{L}{K_{T}}\right)$$

…where $\alpha$ is a proportionality constant.

Day-to-day averaging of experimental split luciferase assay data

In each experimental run (single 96-well plate), it was assumed that luminescence data were proportional to the concentration of complex, up to an additive (background) constant. The background constant was estimated by measuring the luminescence from multiple wells that contained zero target. After background subtraction, the data were assumed to be directly proportional to the concentration of complex. However, the proportionality constant was unknown, and furthermore, could not be assumed to be constant from plate to plate. Therefore, while each experimental run resulted in curves that represented a well-defined relationship between target concentration, $L$, and complex concentration, $C_T$, for a given concentration of probes, $P_{1T}$, $P_{2T}$, curve magnitudes could not be compared plate-to-plate.

To compare data from separate 96-well plates, we assumed these curves were reproducible and only differed by their proportionality constants. Therefore, we adjusted the proportionality constants from each run to minimize the differences between the curves from the set of experimental runs with the same probe concentrations. Once the curves were adjusted, they could be averaged together, and error bars (standard errors) could be found for each value of $L$.

The above procedure worked for a given set of curves as follows. First, each curve in the set was normalized to have a maximum value of 1. Next, the normalized curves were averaged to create an average curve. Then each curve is further normalized by adjusting its proportionality constant so that its least-squares error from the average curve is minimized. After this second normalization step, an updated average curve is calculated, and the process is repeated. This iterative process could proceed until convergence; however, in practice, two iterations (one update to the average curve) was sufficient. The final average curve is referred to as the canonical curve.

This procedure was performed for each set of curves, resulting in canonical curves (with error bars) that represent $C_T$ as proportional to a function of $L_T$ for each value of $P_{1T} = P_{2T}$. The canonical curves were order 1.

Fitting the full model to the data

The full model has five parameters – $K_{\mathrm{D}}^{LB1}, K_{\mathrm{D}}^{LB2}, K_{12}, K_{12L}$, and $K_T$ – and one unknown proportionality constant. However, $K_T$ and the proportionality constant only appear in the equation for $C_T$, and thus can be optimized after solutions for $P_1$, $P_2$, and $L$ are found. Thus, the model fitting was performed in two stages: (1) values of $K_{\mathrm{D}}^{LB1}, K_{\mathrm{D}}^{LB2}, K_{12}, K_{12L}$ are chosen, and the model is solved to find $P_1$, $P_2$, and $L$ for a given $P_{1T}, P_{2T}, L_T$. (2) $K_T$ is varied, and for each value of $K_T$, the best-fit proportionality constant is calculated analytically via linear least squares. The best-fit $K_T$ (for a given choice of $K_{\mathrm{D}}^{LB1}, K_{\mathrm{D}}^{LB2}, K_{12}, K_{12L}$) is the one that minimizes the least squares error.

We used MATLAB's lsqnonlin function to vary $K_{\mathrm{D}}^{LB1}, K_{\mathrm{D}}^{LB2}, K_{12}, K_{12L}$ and find the best-fit values of those parameters. The bounds chosen for $K_{\mathrm{D}}^{LB1}, K_{\mathrm{D}}^{LB2}$ were [0.1,1000] nM. The bounds chosen for $K_{12}, K_{12L}$ were four orders of magnitude above and below their base values of 218 nM and 902 nM$^2$, respectively.

Fitting the reduced model to the data

In the reduced model, we assumed $K_{\mathrm{D}}^{LB1} = K_{\mathrm{D}}^{LB2}$, which reduced the number of model parameters by one. This assumption, together with the two-stage fitting of $K_T$ and $\alpha$, resulted in a 3-parameter problem: fitting $K_{\mathrm{D}}^{LB1}$, $K_{12}$, and $K_{12L}$. The fitting procedure and the upper and lower limits of the parameter search were the same as described above.

Estimation of variance as a function of $C_T$

Throughout the study, both technical and biological replicates were taken. Therefore, both the technical variance, $s_{tech}^2$, and the biological variance, $s_{biol}^2$, could be plotted as functions of the measured output, $C_T$. The total variance, $s_{tot}^2$ is the sum of the two.

Estimation of limit of blank and limit of detection

The limit of blank (LoB) at a given concentration of probes is defined as the lowest value of $L_T$ that is predicted to result in a value of $C_T$ statistically distinguishable from background at an alpha-level of 5%. In this paper, we defined "background" as the value of $C_T$ if no target were present ($M_B$). Thus, the LoB is the value of $L_T$ for which $C_T$ just falls at the edge of the 95% C.I. of the blank measurement:

$$C_T^{LoB} = M_B + 1.645(SD_B)$$

…where $SD_B$ is the standard deviation of the blank measurement as calculated from estimates of $s_{tot}$ (see previous section).

The limit of detection (LoD) at a given concentration of probes is defined as the lowest value of $L_T$ for which the 95% C.I. of the value of $C_T$ does not overlap with the 95% C.I. of the blank measurement:

$$C_T^{LoD} - 1.645(SD_{LoD}) = M_B + 1.645(SD_B)$$

…where $SD_{LoD}$ is the standard deviation of $C_T$ at the LoD, as calculated from estimates of $s_{tot}$ (see previous section).

Estimation of limits of quantification

The limits of quantification (LoQs) are the bounds on an interval in $L_T$ inside of which an unknown value of $L_T$ in an experimental sample may be reliably inferred through a measurement of $C_T$ (at given probe concentrations). In this inverse problem, quantitative inference of $L_T$ can be achieved to a high degree if the slope of $C_T$ vs $L_T$ is large. To that end, in this study, we defined the limits of quantification as where the coefficient of variation ($CV = s_{tot}^L / L_T$) was equal to 0.5, where $s_{tot}^L$ is the estimate of the horizontal error bars of the $C_T$ vs $L_T$ plot. Thus, the quantifiable region is defined as where the CV is less than 0.5.

The horizontal error bars can be found graphically by extending a line segment horizontally from a point on the $C_T$ vs $L_T$ curve until the line segment hits the curves that represent the vertical error bars (see **Fig. 7C**). If the slope of that curve is shallow, then the horizontal error bars can be extended to a length greater than $0.5L_T$, and thus an inference of $L_T$ from a measurement of $C_T$ would be poor.

For given values of total probe concentration, there is an upper LoQ and a lower LoQ. The lower LoQ ($LoQ_L$) will occur where the slope to the left of the $LoQ_L$ becomes too shallow, resulting in a horizontal error bar just larger than $0.5L_T$. The upper LoQ ($LoQ_U$) similarly occurs where the slope to the right of $LoQ_U$ becomes too shallow. This occurs close to where the maximum value of $C_T$ is achieved. It is also possible for the right horizontal error bar to "pass over" the max value of the lower vertical error bar, in which case, the $LoQ_U$ becomes the value of $L_T$ where the right horizontal error bar hits the max value of the lower vertical error bar, regardless of whether the length of that error bar is greater than $0.5L_T$.

### Supplementary Tables

| Gene Fragment | Sequence |
|---|---|
| 1 | CATATGATGTCAATATTAGAAGATCCAGAATTTGTAAAATTAAGACAA<br>TTTAAAGGTAAAGTAAATTTCAATTTAGTTATGCAGATACTGGATGAG<br>ATAGAACTTGATCTAAGGGGAAGTGATAATATCAAGACATCTATAATA<br>TATGTATATTCAAGCCATTTAGATGAGATAAGGAAAAATAAAGAATTC<br>TATGATATGATTGCAGAGATACTACAAAGGTATTACAAAAAAATAGGC<br>ATAGAGAATGTGAATCAGTTGATACTAACTACTATAAAACTCGAG |
| 2 | GTTGGTGGTGGCGGATCAGAAGGAGGCGGTAGCGGGGGCCCTGGTTC<br>GGGAGGGGAAGGTTCTGCTGGGGGAGGGAGCGCTGGCGGGGGGTCTA<br>TGCATATGAAACCTACGGAAAACAACGAAGATTTCAATATCGTGGCCG<br>TGGCTTCAAATTTTGCTACGACGGACCTTGACGCCGACAGAGGAAAGT<br>TACCAGGTAAAAAGCTGCCTTTAGAGGTCCTTAAAGAAATGGAGGCAA<br>ACGCCAGAAAGGCTGGATGTACTAGAGGATGCTTGATCTGCTTATCTC<br>ACATCAAATGTACTCCAAAAATGAAGAAATTTATCCCTGGCAGATGCC<br>ACACCTACGAGGGTGACAAGGAGAGCGCTCAGGGTGGAATTGGAGAA<br>GCTATCGTCGACATCCCTGAAATTCCAGGCTTCAAGGACTTAGAGCCT<br>ATGGAGCAATTTATCGCACAGGTCGACCTTTGTGTTGACTGCACCACC<br>GGATGCCTTAAGGGTCTGGCAAACGTCCAGTGTTCCGACCTTCTGAAG<br>AAATGGCTGCCACAGAGATGTGCGACGTTTGCAAGCAAGATTCAAGGC<br>CAAGTCGATAAGATCAAGGGCGCAGGAGGAGATACTAGTGGCGCTGG<br>CGGAAGCCCTGGGGGCGGGAGCGGTGGCTCTGGTTCTTCTGCTAGTGG<br>CGGCTCAACATCT |
| 3 | GTTGGTGGTGGCGGATCAGAAGGAGGCGGTAGCGGGGGCCCTGGTTC<br>GGGAGGGGAAGGTTCTGCTGGGGGAGGGAGCGCTGGCGGGGGGTCTA<br>TGCATATGGTTTTCACCTTAGAGGATTTCGTAGGCGACTGGAGACAGA<br>CGGCAGGCTATAACTTGGACCAAGTGCTGGAACAGGGCGGTGTCAGCT<br>CCTTATTCCAGAATCTGGGGGTATCCGTTACTCCTATACAGAGAATAGT<br>TCTTTCCGGAGAAAACGGGCTAAAGATAGATATTCATGTCATTATTCCT<br>TACGAAGGATTGTCCGGCGACCAAATGGGTCAAATCGAAAAGATTTTC<br>AAGGTGGTTTACCCGGTAGACGATCATCACTTCAAGGTGATTTTGCATT<br>ACGGTACCTTAGTTATTGACGGGGTAACGCCAAACATGATAGACTACT<br>TTGGCAGACCCTACGAGGGAATTGCAGTTTTCGATGGTAAAAAGATCA<br>CGGTTACGGGTACTCTGTGGAACGGGAATAAAATTATAGATGAGCGTT<br>TAATCAATCCAGATGGATCACTACTTTTTAGAGTGACAATTAACGGTGT<br>CACTGGCTGGAGGTTATGCGAAAGAATACTTGCGACTAGTGGCGCTGG<br>CGGAAGCCCTGGGGGCGGGAGCGGTGGCTCTGGTTCTTCTGCTAGTGG<br>CGGCTCAACATCT |
| 4 | CAGACGGAGATATGCATATGCACCATCACCATCACCATGCTAGCATGG<br>CGACCGTGAAATTTAAATATAAAGGCGAAGAAAAACAGGTGGATATT<br>AGCAAAATTGGTGCTGTGGCGCGCCTAGGCAAATATATTATTTTTGTGT<br>ATGATCTGGGCGGCGGCAAAACAGGCTTGGGCCTAGTGAGCGAAAAA<br>GATGCGCCGAAAGAACTGCTGCAGATGCTGGAAAAACAGAAAAAAGG<br>ATCCGTTGGTGGTGGCGGATCAGAAGGAGGCGGTAGCGGGGGCCCTG<br>GTTCGGGAGGGGAAGGTTCTGCTGGGGGAGGGAGCGCTGGCGGGGGG |

| | |
|---|---|
| | TCTATGGTTTTCACCTTAGAGGATTTCGTTGGAGACTGGGAGCAAACA<br>GCCGCCTATAATTTGGACCAGGTCCTGGAGCAGGGGGGAGTCTCTAGC<br>TTGTTGCAAAATTTAGCAGTGAGTGTTACGCCCATCCAGCGCATTGTAC<br>GCTCAGGGGAGAACGCTCTGAAAATTGATATTCACGTTATCATCCCCT<br>ACGAGGGGCTTAGCGCGGATCAAATGGCTCAGATTGAGGAGGTTTTTA<br>AAGTCGTTTATCCAGTGGACGACCATCATTTTAAAGTGATTCTGCCATA<br>CGGCACCTTGGTGATTGACGGCGTAACCCCTAACATGCTGAATTACTTC<br>GGTCGTCCCTATGAGGGTATTGCTGTGTTCGACGGAAAAAAGATCACT<br>GTAACTGGAACTCTTTGGAATGGAAACAAAATCATCGACGAACGCCTT<br>ATCACGCCCGACGGTTCGATGTTGTTCCGTGTGACAATTAACAGT |
| 5 | ATGGTTTTCACCTTAGAGGATTTCGTTGGAGACTGGGAGCAAACAGCC<br>GCCTATAATTTGGACCAGGTCCTGGAGCAGGGGGGAGTCTCTAGCTTG<br>TTGCAAAATTTAGCAGTGAGTGTTACGCCCATCCAGCGCATTGTACGCT<br>CAGGGGAGAACGCTCTGAAAATTGATATTCACGTTATCATCCCCTACG<br>AGGGGCTTAGCGCGGATCAAATGGCTCAGATTGAGGAGGTTTTTAAAG<br>TCGTTTATCCAGTGGACGACCATCATTTTAAAGTGATTCTGCCATACGG<br>CACCTTGGTGATTGACGGCGTAACCCCTAACATGCTGAATTACTTCGGT<br>CGTCCCTATGAGGGTATTGCTGTGTTCGACGGAAAAAAGATCACTGTA<br>ACTGGAACTCTTTGGAATGGAAACAAAATCATCGACGAACGCCTTATC<br>ACGCCCGACGGTTCGATGTTGTTCCGTGTGACAATTAACAGTGTCACC<br>GGATACCGTCTTTTTGAAGAAATCCTGACTAGTGGCGCTGGCGGAAGC<br>CCTGGGGGCGGGAGCGGTGGCTCTGGTTCTTCTGCTAGTGGCGGCTCA<br>ACATCTCCTAGGATGGCGACCGTGAAATTTAAATATAAAGGCGAAGAA<br>AAACAGGTGGATATTAGCAAAATTTGGAAGGTGTGGCGCGAAGGCAA<br>AATTATTACTTTTACGTATGATCTGGGCGGCGGCAAAGCAGGCTTAGG<br>CTTTGTGAGCGAAAAAGATGCGCCGAAAGAACTGCTGCAGATGCTGGA<br>AAAACAGAAAAAA |
| 6 | ATGGCGACCGTGAAATTTAAATATAAAGGCGAAGAAAAACAGGTGGA<br>TATTAGCAAAATTCTCTTCGTGCTACGCTTTGGCAAAGACATTTTTTTTT<br>GGTATGATCTGGGCGGCGGCAAACATGGCGTGGGCGAGGTGAGCGAA<br>AAAGATGCGCCGAAAGAACTGCTGCAGATGCTGGAAAAACAGAAAAA<br>ACCTAGGGTTGGTGGTGGCGGATCAGAAGGAGGCGGTAGCGGGGGCC<br>CTGGTTCGGGAGGGGAAGGTTCTGCTGGGGGAGGGAGCGCTGGCGGG<br>GGGTCTATGGTTTTCACCTTAGAGGATTTCGTTGGAGACTGGGAGCAA<br>ACAGCCGCCTATAATTTGGACCAGGTCCTGGAGCAGGGGGGAGTCTCT<br>AGCTTGTTGCAAAATTTAGCAGTGAGTGTTACGCCCATCCAGCGCATT<br>GTACGCTCAGGGGAGAACGCTCTGAAAATTGATATTCACGTTATCATC<br>CCCTACGAGGGGCTTAGCGCGGATCAAATGGCTCAGATTGAGGAGGTT<br>TTTAAAGTCGTTTATCCAGTGGACGACCATCATTTTAAAGTGATTCTGC<br>CATACGGCACCTTGGTGATTGACGGCGTAACCCCTAACATGCTGAATT<br>ACTTCGGTCGTCCCTATGAGGGTATTGCTGTGTTCGACGGAAAAAAGA<br>TCACTGTAACTGGAACTCTTTGGAATGGAAACAAAATCATCGACGAAC<br>GCCTTATCACGCCCGACGGTTCGATGTTGTTCCGTGTGACAATTAACAG<br>T |

**Table S1.** Gene Fragments used to construct plasmids containing Sso6904 or split luciferase probes.

| Primer | Sequence |
|---|---|
| Pf1 | GATGCACATATGATGGCGACCGTGAAATTTAAATAT |
| Pr1 | CCGCCGCTCGAGTTTTTTCTGTTTTTCCAGCATCTG |
| Pf2 | CCCGGGGTTGGTGGTGGCGGATCAG |
| Pr2 | CCTAGGAGATGTTGAGCCGCCACTAG |
| Pf3 | ATGCATATGGTTTTCACCTTAGAGGATTTC |
| Pr3 | ACTAGTCGCAAGTATTCTTTCGCATAACC |
| Pf4 | GAGTACCCATGGGCAGCAGCCATCACCATCACCATCACGCTAGCGGCA GTATGGCGACCGTGAAATTTAAATAT |
| Pr4 | CTCGAGTTATCAGTTTTCTCCGGAAAGAACTATTC |
| Pf5 | CATATGGGGCTAAAGATAGATATTCATGTC |
| Pr5 | GAGTCACTCGAGTTTTTTCTGTTTTTCCAGCATCTG |
| Pf6 | CAGACGGAGATATGCATATGCAC |
| Pr6 | GCTCGTCTCGAGTTATCAACTGTTAATTGTCACACGGAACAA |
| Pf7 | GAGTCAGCTAGCATGGCGACCGTGAAATTTAAATAT |
| Pr7 | GAGTCAGGATCCTTTTTTCTGTTTTTCCAGCATCTG |
| Pf8 | CATATGGTCACCGGATACCGTCTTTTTG |
| Pf9 | GCTTAGCATATGATGGCGACCGTGAAATTTAAA |
| Pr9 | GCTTAGCTCGAGACTGTTAATTGTCACACGGAA |
| Pf10 | ATTCGACCTAGGATGGCGACCGTGAAATTTAAA |
| Pr10 | ATTCGACTCGAGTTTTTTCTGTTTTTCCAGCATCT |

**Table S2.** Primers used to construct plasmids containing Snof3 or split luciferase probes. Sequences are given from 5’ to 3’.

### Supplementary Figures

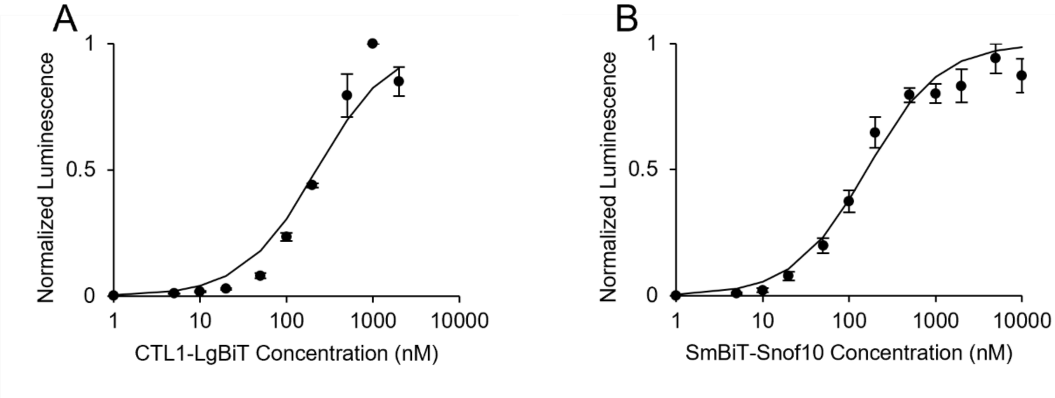


**Figure S1.** Apparent affinity between the lysozyme detection probes CTL1-LgBiT and SmBiT-NTL1 (A) and between the Sso6904 detection probes Snof3-LgBiT and SmBiT-Snof10 (B) were estimated by conducting mix-and-read assays with a constant concentration of one probe (SmBiT-NTL1 or Snof3-LgBiT), varying amounts of the other probe (CTL1-LgBiT or SmBiT-Snof10), and no target protein. Luminescence was normalized by the maximum luminescence of each repeat and affinity was calculated using a global non-linear least squares fit across three independent replicates for each set of probes. The apparent affinity between CTL1-LgBiT and SmBiT-NTL1 is 210 nM (68% confidence interval: 160 nM-271 nM) and the apparent affinity between Snof3-LgBiT and SmBiT-Snof10 is 150 nM (68% confidence interval: 114 nM-191 nM). Error bars correspond to standard error.

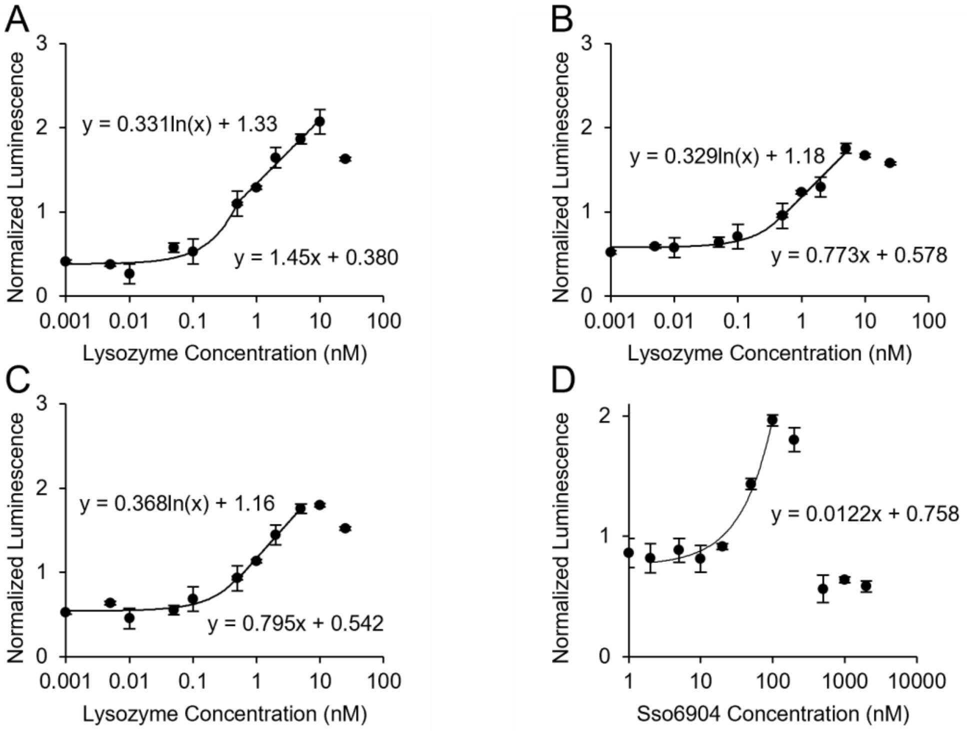


**Figure S2.** (A-C) Linear and logarithmic ranges of split luciferase lysozyme detection system with 1 nM (A), 5 nM (B), or 10 nM (C) probes. (D) Linear range of split luciferase Sso6904 detection system with 100 nM probes. All data are shown on semi-log plots.

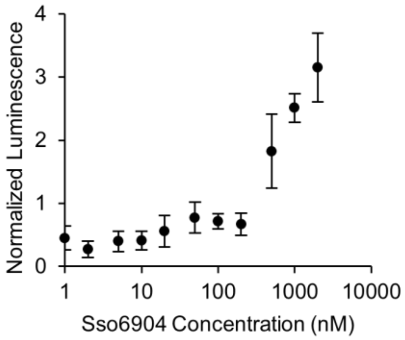


**Figure S3.** Split luciferase Sso6904 detection assay. Detection of Sso6904 at different concentrations was measured with the mix-and-read split luciferase detection assay using 10 nM Snof3-LgBiT and 10 nM SmBiT-Snof10. Equilibration step and detection incubation times were 4 hours and 1 hour, respectively—times optimized for the lysozyme detection system. Luminescence is normalized by the mean luminescent signal at all Sso6904 concentrations for each repeat. Three independent replicates were conducted. Error bars indicate standard error.
